# ExMODE: A Multi-Omics Repository for Extremophile Adaptation and Bioprospecting

**DOI:** 10.64898/2026.04.27.720953

**Authors:** Denghui Li, Kailong Ma, Yulu Zhang, Jieni Wang, Zhen Cui, Xiaoqiang Li, Weiwen Wang, Jiawei Tong, Yang Guo, Zongan Wang, Peiyun Zeng, Jieyu Wang, Xiaokai Xu, Nannan Zhang, Yingying Zhang, Jing Chen, Qingjiang Hu, Wenzhen Yang, Zhiyong Li, Tao Yang, Wensi Du, Zhicheng Xu, Zhen Yue, Jian Wang, Guangyi Fan, Weijia Zhang, Xun Xu, Liujie Huo, Xiaofeng Wei, Liang Meng, Shanshan Liu

**Affiliations:** BGI Research, Sanya 572025, China; BGI Research, Shenzhen 518083, China; BGI Research, Wuhan 430074, China; State Key Laboratory of Microbial Technology, Shandong University, Qingdao 266237, China; Institute of Oceanology, Chinese Academy of Sciences, Qingdao 266071, China; BGI, Shenzhen 518083, China; BGI Research, Qingdao 266555, China; Institute of Deep-sea Science and Engineering, Chinese Academy of Sciences, Sanya 572000, China; Hainan Technology Innovation Center for Marine Biological Resources Utilization(Preparatory Period), BGI Research, Sanya 572025, China; Shenzhen Key Laboratory of Bioenergy, BGI Research, Shenzhen 518083, China; State Key Laboratory of Genome and Multi-omics Technologies, BGI Research, Shenzhen 518083, China; Institution of Deep-Sea Life Sciences, IDSSE-BGI, Hainan Deep-sea Technology Laboratory, Sanya 572000, China; Shenzhen Key Laboratory of Marine Biology Genomics, BGI Research, Shenzhen 518083, China

## Abstract

Extreme environments, though hostile to most life forms, host specialized extremophile communities that have redefined biological cognition and emerged as vital biotechnological resources, with their unique adaptive traits and bioactive molecules driving advances in multiple scientific and industrial fields. However, research on extremophiles is hindered by limitations in culture-based methods, fragmented multi-omics data with non-uniform annotation standards across repositories, the lack of cross-extreme comparative research in existing resources, and the singularity of data dimensionality that neglects key structural information, all of which restrict the functional interpretation of extremophile microbes and the exploitation of their bioprospecting potential. To tackle these challenges, we developed ExMODE (https://db.genomics.cn/exmode/), a comprehensive multi-omics database platform dedicated to extremophiles. It centrally integrates multi-omics data from diverse extreme habitats with a standardized annotation framework, resolving data fragmentation and enabling systematic cross-environment comparative analyses to elucidate extremophile adaptive mechanisms. Moreover, ExMODE aggregates multi-dimensional datasets including genes, genomes, secondary metabolite sequences and protein structures, overcoming the constraints of single-dimensional data and significantly improving the efficiency of biotechnological resource discovery from extreme microorganisms.

## INTRODUCTION

While conditions such as extreme thermal gradients, hypersalinity, and intense hydrostatic pressure render most habitats uninhabitable, these specific stressors define what are termed extreme environments (1). Rather than being sterile wastelands, habitats like glaciers, hydrothermal vents and acid mine drainage foster specialized biological communities. Within these environments, extremophiles (e.g., thermophiles, psychrophiles and acidophiles) do not merely survive but exploit these rigorous conditions as ecological niches (1-4).

The exploration of extremophiles has reshaped our understanding of biological limits and diversity (5,6). Beyond elucidating key insights of biomolecular stability, evolutionary origins, and ecosystem resilience (4,7-10). these organisms represent a substantial reservoir of biotechnological resources. Extremophiles yield robust enzymes and bioactive compounds with functional versatility in industrial catalysis and drug discovery (3,4,11,12). This dual significance is best exemplified by the ubiquitous Taq polymerase, which widespread use in molecular biology (13), and the recent discovery of deep-sea polyethylene terephthalate (PET) hydrolases that underscore the immense bioprospecting potential of extreme habitats (14).

Despite these advances, significant impediments persist. Culture-based approaches recover only a limited fraction of microbial diversity, as many taxa remain recalcitrant to laboratory cultivation (15,16). While metagenomics bypasses the need for cultivation, the resulting data are often stored across multiple repositories with heterogeneous annotation standards, making it difficult to integrate them effectively into a unified multi-omics framework (17,18). Furthermore, many existing resources are primarily concentrated on generalized habitats or specific extreme environments, thereby lacking the cross-extreme comparative depth necessary to decouple convergent evolutionary traits from niche-specific adaptive mechanisms. Crucially, current repositories frequently suffer from a singularity in data dimensionality, focusing on gene sequences while lacking critical 3D structural information. Such dimensional disparities impose substantial constraints on the functional interpretation of microbial dark matter, leaving the considerable bioprospecting potential of extreme ecosystems largely unexploited..

To address this, we developed ExMODE (https://db.genomics.cn/exmode/), a comprehensive platform that centrally integrates multi-omics data derived from diverse extreme environments. By implementing a rigorous and standardized annotation framework, ExMODE effectively mitigates data fragmentation and inconsistencies in annotation, while enabling systematic cross-environment comparative analyses. This foundation provides robust support for elucidating adaptive mechanisms in extremophiles. In addition, the platform systematically aggregates large-scale datasets encompassing genes, genomes, secondary metabolite sequences, and protein structures, thereby overcoming the limitations of single-dimensional data and enhancing the efficiency of biotechnological resource discovery from extreme microorganisms.

## RESULTS

### Data summary

A total of 3,518 samples were collected globally, derived from 1,045 projects, with a cumulative data volume exceeding 130 TB. Following manual curation, samples were categorized into five extreme habitat types: Thermal (769), Cryogenic (406), Acidic (150), Saline (502), and Hyperbaric (2,177) (Table S1, Methods); some polyextremophilic samples were assigned to multiple categories. Using standardized analytical pipelines, we generated 1.35 billion non-redundant genes (unigenes), 5.25 million representative protein structures, 67,026 metagenome-assembled genomes (MAGs), and 164,132 biosynthetic gene clusters (BGCs) (Figure 1).

**Figure 1.**
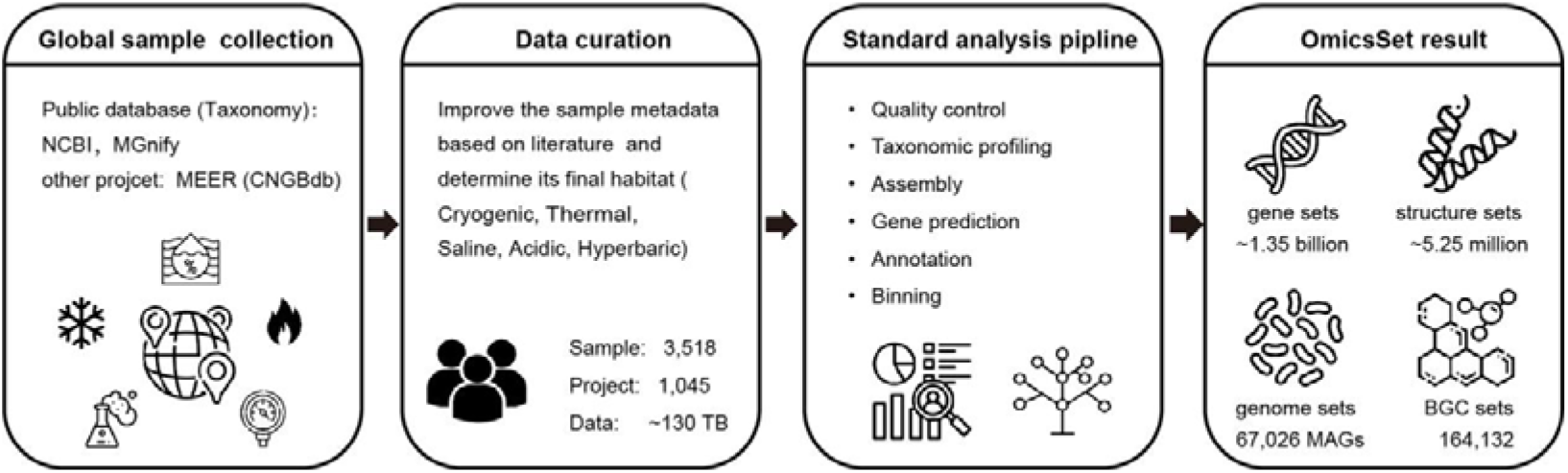
The ExMODE data processing pipeline

### Gene sets and Structure sets

Based on a 95% sequence similarity threshold, non-redundant gene sets were constructed for distinct habitats: Thermal (324,605,762 unigenes), Cryogenic (170,227,909 unigenes), Acidic (89,775,476 unigenes), Saline (237,053,808 unigenes), and Hyperbaric (526,088,471 unigenes) (Figure 2A). Each set was systematically annotated against the CAZy, KEGG, eggNOG, and CARD databases (19-22). Except for CARD annotations, the majority of genes lacked precise functional assignments, indicating a vast reservoir of uncharacterized genetic “dark matter” with significant bioprospecting potential (23-25). To facilitate resource discovery and utilization, we predicted representative protein structures from each gene set, identifying 1,097,941 (Thermal), 568,397 (Cryogenic), 325,033 (Acidic), 774,228 (Saline), and 2,485,847 (Hyperbaric) structures (Methods). These representative structures accounted for at least 40% of the unigenes in each habitat (Figure S1). The predicted structures were assigned to three confidence categories (high, good, and low), with more than 55% in each habitat being of medium or high quality, facilitating user structure retrieval based on diverse scenarios (Figure 2B).

**Figure 2.**
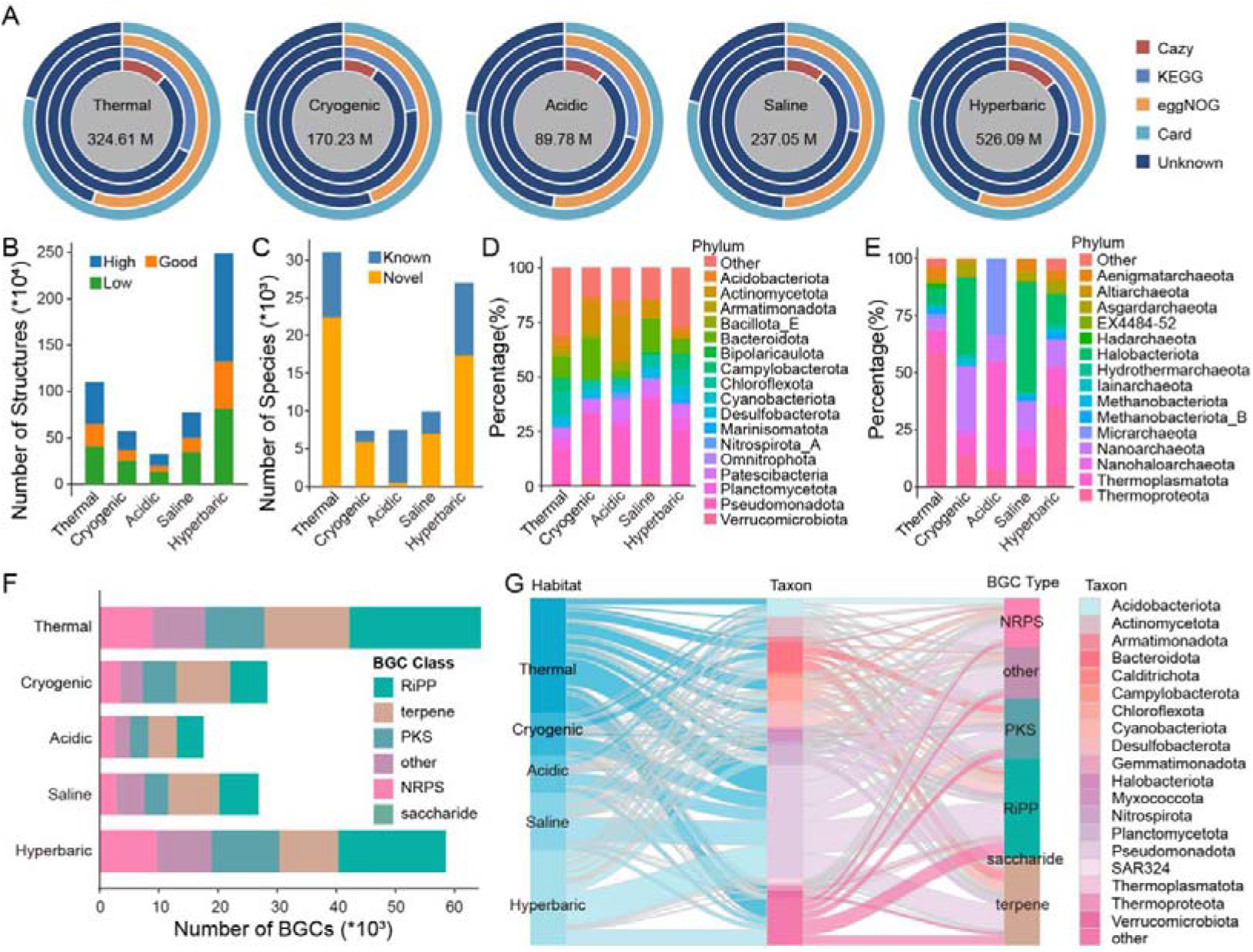
The Results of omics analysis. (A) Functional annotations of gene sets from different extreme habitats. (B) Structure sets of different extreme habitats. High: high-confidence (pLDDT > 0.7 and pTM > 0.7). Good: good-confidence (pLDDT > 0.5 and pTM > 0.5). Low: low-confidence (pLDDT < 0.5 or pTM < 0.5). (C) The taxonomic annotations (species level) of MAGs from different extreme habitats. (D) Taxonomic breakdown of the top 10 dominant bacterial phyla in genome sets from different extreme habitats. Phyla not in the top 10 are collectively referred to as “Other”. (E) Taxonomic breakdown of the top 10 dominant archaeal phyla in genome sets from different extreme habitats. Phyla not in the top 10 are collectively referred to as “Other”. (F) Number of BGCs in different extreme environments. (G) Associations between phylum-level taxa, BGC types, and habitats in different extreme environments. Phyla harboring fewer than 1000 BGCs are grouped as “Other”.

### Genome sets

We assembled habitat-specific genomic datasets comprising 31,032 Thermal, 7,347 Cryogenic, 7,442 Acidic, 9,913 Saline, and 26,983 Hyperbaric MAGs. Analysis revealed a high proportion of novel species-level lineages across all environments except acidic habitats, highlighting an expansive reservoir of untapped microbial diversity (Figure 2C). While dominant bacterial phyla remained relatively consistent across habitats (Figure 2D), archaeal distributions showed clear habitat-specific distribution patterns: Halobacteriota dominated saline sites, whereas Thermoproteota were most prevalent in thermal environments (Figure 2E). These datasets provide a foundational resource for investigating microbial adaptation and bioprospecting within extreme niches.

### BGC sets

BGCs encode a diverse array of secondary metabolites—such as antibiotics, antifungals, and anticancer agents—that mediate microbial interaction and environmental adaptation. While BGCs from common habitats have been extensively studied, those from extreme environments remain underexplored and hold great potential for yielding novel chemical scaffolds with unique bioactivities. To systematically profile this biosynthetic potential, we constructed BGC sets using antiSMASH (26), identifying over 164,000 BGCs across extreme habitats, including 64,779 Thermal, 28,357 Cryogenic, 17,574 Acidic, 26,879 Saline, and 58,519 Hyperbaric BGCs. Comparative analysis revealed distinct BGC distribution patterns: terpenes dominated Cryogenic, Acidic, and Saline environments, mirroring their prevalence in soil and marine systems (27,28), while Ribosomally synthesized and Post-translationally modified Peptides (RiPPs) were most abundant in Thermal and Hyperbaric habitats (Figure 2F). Cross-habitat association analysis further highlighted Pseudomonadota, Bacteroidota, Chloroflexota, and Actinomycetota as major BGC-producing phyla (Figure 2G). Notably, the candidate genus *UBA10656* in Thermal habitats and *Rhodococcus* in Hyperbaric habitats exhibited exceptional biosynthetic density (avg.≥20 BGCs per MAG, Table S2). These findings underscored extreme environments as rich reservoirs of untapped biosynthetic diversity, with clear ecological and taxonomic specialization in secondary metabolite production.

### Web interface

ExMODE was developed to facilitate exploration and utility of multi-omics data from extreme environments. By integrating seven specialized modules—MapFinder, SeqFinder, StructFinder, Browse, Omicset, Collection, and Tools— the platform offers a versatile framework for data retrieval and analysis. MapFinder enables geospatial visualization, allowing users to interrogate sample distributions directly via an interactive map (Figure 3B). For sequence analysis, SeqFinder expedites the retrieval of genes, genomes, and biosynthetic gene clusters (BGCs), delivering associated sequences and metadata (Figures 3C). Complementing this, StructFinder performs structure-based similarity searches on 3D protein models to pinpoint putative functional homologs (Figures 3D). Users can also navigate the data hierarchically via the Browse interface (Figure 3E). For systematic exploration, the Omicset module organizes fragmented data into non-redundant gene, genome, and BGC catalogs, providing a standardized resource for investigating microbial diversity and biosynthetic potential (Figure 3F). The Collection module serves as a repository of curated resources from published key studies in extreme environments, providing a dedicated foundation for further research (Figure 3G). Finally, the Tools module incorporates mature utilities, including AlphaFold3 (29), antiSMASH (26), and eggNOG-mapper (30), to enable customized downstream analyses (Figure 3H).

**Figure 3.**
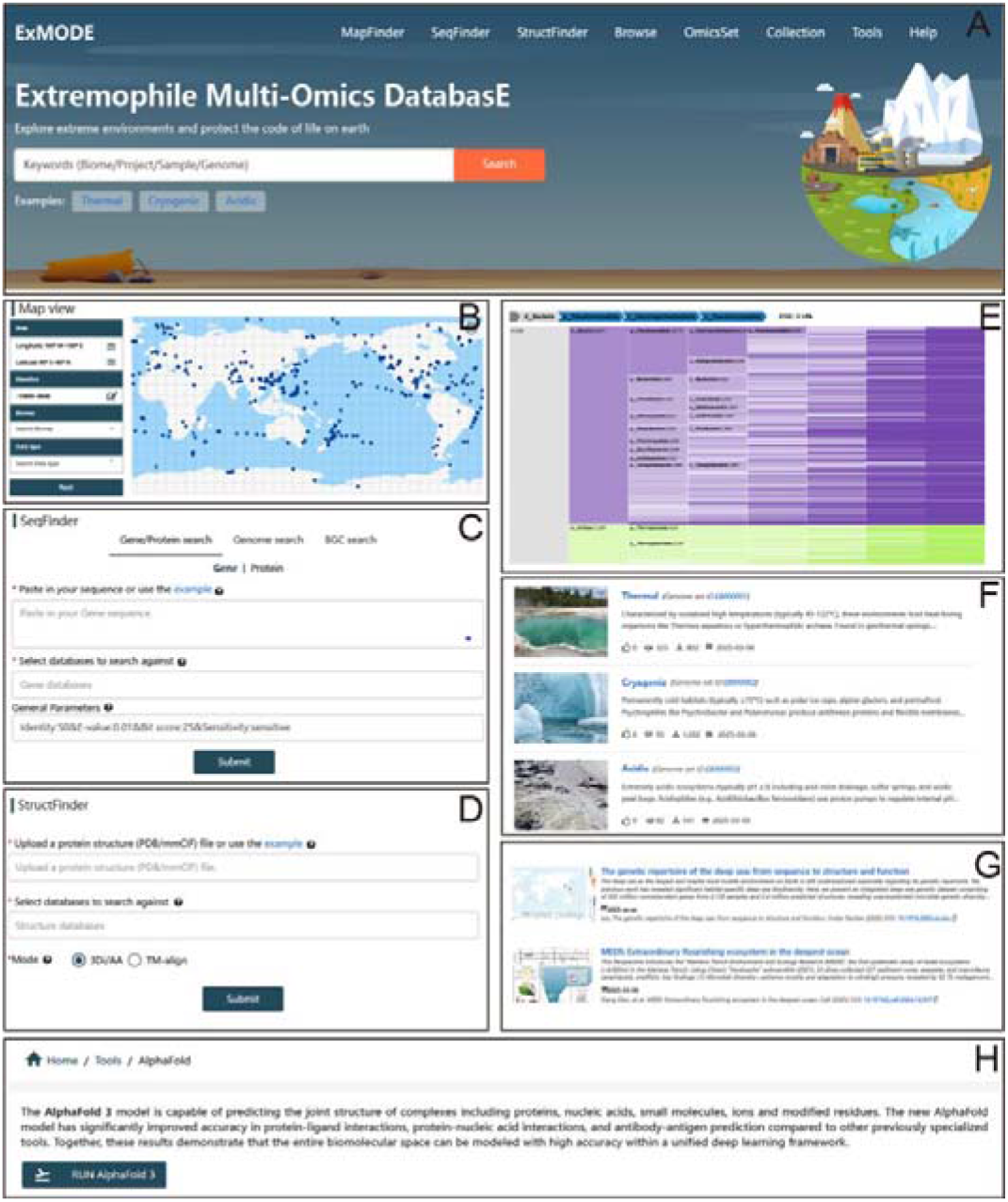
The ExMODE web server. (A) Homepage Interface. (B) Mapfinder Interface. (C) SeqFinder Interface. (D) StructFinder Interface. (E) Browse Interface. (F) OmicSet Interface. (G) Collection Interface. (H) Tools Interface.

### Comparison of major microbial and multi-omics databases

We compared ExMODE with general metagenomic databases (MGnify (31), gcMeta (32)) and specialized extreme-environment resources (4GDB (33), DOO (34)) to evaluate its performance in resource localization and functional exploration (Table 1). Existing databases differ in scope and data composition but remain limited in multi-omics integration and functional mining for extreme environments.

**Table 1.** Overview of database features and capabilities across ExMODE and existing resources.

| Category | ExMODE | MGnify | gcMeta | 4GDB | DOO |
| --- | --- | --- | --- | --- | --- |

| Research Focus | Extremophiles | Global microorganisms | Global microorganisms | Psychrophiles | Deep ocean organisms |
| --- | --- | --- | --- | --- | --- |
| Ecological Coverage | Thermal, Cryogenic, Acidic, Saline and Hyperbaric | ~300 biomes (human, marine, soil, etc.) | ~50 biomes (human, marine, soil, etc.) | Cryogenic | Deep-sea |
| Latest update | This work | 2023 | 2025 | 2025 | 2025 |
| Data Scale<sup>a</sup> | 3518 | 343695 | 104266 | 3262 | 2149<sup>b</sup> |
| Data Types | Metatranscriptome and Metagenome | Amplicon, Metatranscriptome and Metagenome | Metagenome | Amplicon, Genome and Metagenome | Genome, Transcriptome and Metagenome |
| Test search | ✓ | ✓ | ✓ | ✓ | ✗ |
| Gene/Protein search | ✓ | ✓ | ✗ | ✓ | ✓ |
| Genome search | ✓ | ✓ | ✗ | ✗ | ✗ |
| BGC search | ✓ | ✗ | ✗ | ✗ | ✗ |
| Structure search | ✓ | ✗ | ✗ | ✗ | ✗ |
| Resource set | Protein, Genome and BGC | Protein and Genome | Genome | Gene and Genome | No |

<sup>a</sup> The number of samples contained in the database.
<sup>b</sup> Data from 68 species, encompassing 72 genomes, 950 bulk transcriptomes, 15 single-cell transcriptomes, and 1112 metagenomes

MGnify (31) and gcMeta (32) provide broad ecosystem coverage but are dominated by data from conventional habitats, with relatively sparse representation of extreme environments. Although MGnify encompasses extensive multi-omics datasets, a significant proportion is derived from amplicon sequencing, which compared with metagenomic data, provides more limited resolution for genome-resolved and functional analyses. In contrast, 4GDB (33) and DOO (34) focus on specific extreme settings but are restricted in ecological diversity and data structure. 4GDB (33) primarily targets low-temperature environments, whereas DOO (34) largely relies on host-associated metagenomic data, limiting its representation of natural deep-sea microbial communities. By contrast, ExMODE integrates multi-omics data across five major extreme environments and incorporates non-redundant genes, high-quality MAGs, and BGCs, forming one of the most comprehensive resources currently available. It further enables multi-level retrieval from genes to genomes and BGCs, supporting cross-scale functional analysis. Notably, ExMODE uniquely supports protein structure similarity search, addressing the low sequence conservation yet conserved structural features typical of extremophile proteins, and improving the detection of distant homologs.

Collectively, ExMODE expands both the scale and analytical depth of extreme-environment resources, providing an effective platform for functional discovery and biotechnological exploration.

## DISCUSSION AND CONCLUSION

The ExMODE database, which constitutes a systematic integration of multi-omics data from extreme habitats, specifically Thermal, Cryogenic, Acidic, Saline, and Hyperbaric systems is designed to resolve the microbial genomic and functional landscape of these underexplored biomes as well as to unlock their biotechnological potential. In contrast to general repositories such as MGnify (31) and IMG/M (35), which provide broad taxonomic overviews but lack environmental context and curated annotations, ExMODE establishes a dedicated framework that explicitly links microbial diversity with environmental metadata. This integrated framework not only enables rigorous inquiry into the adaptive mechanisms and ecological interactions that underpin survival in extreme conditions but also accelerates the discovery of novel biomolecules and the organisms that produce them.

Our analyses reveal distinctive genomic and metabolic features of extreme environments compared to conventional biomes. A notably higher proportion of genes of unknown function underscores extreme habitats as a major reservoir of genomic “dark matter” (24). This extensive uncharacterized diversity, coupled with the prevalence of novel species-level lineages across most extreme habitats, reflects strong environmental filtering and niche-driven specialization. Furthermore, we observed clear habitat-specific patterns in archaeal phylum-level distribution and distinct biogeographic signatures in BGCs— such as the dominance of RiPPs in Thermal and Hyperbaric environments—suggesting that extreme physicochemical conditions drive divergent evolutionary trajectories in secondary metabolism. These findings collectively position extreme environments as unique hotspots for discovering novel enzymes, antimicrobial agents, and bioactive compounds, with profound implications for both understanding adaptive evolution and advancing biotechnological innovation.

However, several limitations remain. The pervasive use of short-read sequencing technologies often fractures valuable BGCs and complicates their functional characterization, which impedes the reconstruction of complete biosynthetic pathways and large genomic elements. To overcome this, future updates of ExMODE will prioritize the incorporation of long-read sequencing data to enhance contig contiguity and facilitate more accurate metabolic inference.

In summary, ExMODE is of twofold value by centralizing access to multi-omics data on extremophiles: (1) enabling unprecedented insight into microbial adaptation while creating a structured repository for testing hypotheses regarding biosynthetic innovation and evolutionary trajectory; (2) streamlining the pipeline from gene discovery to application, holding immediate promise for the development of novel enzymes, antimicrobials, biomaterials, and other high-value bio-products. We envision ExMODE as a dynamic, community-driven resource that will continuously assimilate new data to advance the understanding of extremophile evolution and to support biotechnology innovation.

## METHODS

### Data collection

We conducted a comprehensive survey of published datasets on extremophiles from public repositories, including NCBI, MGnify, and CNGBdb. For each sample, we collected both the raw metagenomic/metatranscriptomic sequencing data and the associated metadata. This metadata was then used to systematically categorize the samples into five distinct extreme environments based on taxonomic assignments and habitat descriptors. The specific classification criteria were as follows: Thermal environment primarily comprised samples from hot spring and hydrothermal vent; Cryogenic environment included glacier, permafrost, and polar lake samples; Saline environment was represented by salt marsh and salt lake samples; Acidic environment mainly consisted of acid mine drainage samples; and Hyperbaric environment was defined by deep-sea metagenomic samples from depths exceeding 1000 meters (>10 MPa). Following this classification and a rigorous quality assessment, we ultimately retained 3,518 samples with paired-end libraries, thereby constructing a more complete and reliable extremophiles dataset. Our hyperbaric environment incorporates the deep-sea metagenomic resources described by Guo et al. (36) as a core dataset. This includes the systematic integration of 502 million nonredundant genes and 2.4 million protein structures from 2,138 hyperbaric samples into the ExMODE multi-omics framework.

### Filtering, taxonomic classification and assembling

To address the challenges of data heterogeneity, we have developed a standardized analytical pipeline. SRA-formatted files were converted to FASTQ format using the SRA Toolkit’s fastq-dump (v2.10.8, https://github.com/ncbi/sra-tools). Raw sequencing data were filtered using fastp (v0.20.1) (37) with default parameters. Taxonomic classification was conducted using MetaPhlAn4 (v4.1.0) (38) with default parameters, and community structures were calculated. Clean reads were assembled using MEGAHIT (v1.2.9) (39) with parameters “--presets meta-sensitive”.

### Construction of datasets of protein sequences and structures

For assembly-based analysis, protein-coding genes were predicted from contigs of each sample using MetaGeneMark (v3.38) (40) with default parameters. Predicted protein sequences from multiple samples within the same sub-environment were deduplicated via MMseqs2 (v7e28) (41) with parameters “--min-seq-id 0.95 -c 0.90”, generating a non-redundant gene set. Following initial deduplication, the non-redundant gene sets from distinct extreme habitats were further clustered separately using MMseqs2 (v7e28) (41) with the specified parameters “--min-seq-id 0.20 -c 0.50”. Large protein families derived from gene sets (> 30 members) were subsequently utilized to construct protein structure sets. For each cluster, all sequences identified as complete open reading frames (ORFs) were aligned to the centroid sequence employing BLASTP (v2.16.0) (42). The top two centroid-matching sequences from each cluster were processed with ESMFold (43) under default parameters for protein structure prediction. The predicted structure exhibiting the higher pTM score within each cluster was designated as the representative and incorporated into the protein structure sets. Functional annotation of coding genes was performed using eggNOG-mapper (v2.1.9) (30) by mapping to the eggNOG database (v5.0) (21). This approach assigned protein sequences to orthologous groups, providing functional annotations and enabling KEGG pathway inference (20). For specialized functional annotation, Carbohydrate-Active enZYmes (CAZy) (19) were identified using DIAMOND (2.1.6) (44), while antibiotic resistance genes were detected through the Comprehensive Antibiotic Resistance Database (CARD) using Resistance Gene Identifier (RGI) (v6.0.3) with parameters “--local --clean --include_loose -t protein” (22).

### Metagenomic binning

For metagenomic binning analysis, metagenome-assembled genomes were retrieved using the “binning” modules of MetaWRAP (v1.3) (45) with parameters “--metabat2 --concoct --maxbin2 -l 1000”. Incomplete genomes were filtered based on the lineage_wf workflow: the completeness >50% and contamination <10% were retained. To remove redundant genomes, MAGs from multiple samples in the same sub-environments were dereplicated using dRep (v3.3.0) (46) with an average nucleotide identity (ANI) threshold of 95% and at least 30% overlap between genomes (parameters: --completeness 50 --contamination 10 --P_ani 0.9 --S_ani 0.95 --cov_thresh 0.3). All MAGs were taxonomically classified using GTDB-Tk (v2.3.2) (47) with reference to the Genome Taxonomy Database (GTDB r214) (48), yielding standardized taxonomic classifications with parameters “classify_wf --skip_ani_screen “.

To obtain the protein sequences of each MAG, gene open reading frames (ORFs) for the MAGs were predicted using Prodigal (v2.6.3) (49). Subsequently, these protein sequences were functionally annotated by comparing them with the COG and KEGG databases (20,50) using DIAMOND (2.1.6) (44).

### Biosynthetic gene cluster analysis

Concurrently, BGCs within MAGs were identified using antiSMASH (v7.1.0) (26) with the following parameters: “-genefinding-tool prodigal, -cb-knownclusters”. Clustering analysis was performed using BiG-SCAPE (v 2.0.0b8) (51) with the PFAM database (v37.0) (52). The analysis utilized antiSMASH-generated GenBank files (.gbk) as input, executed with the parameters: “--mix --hybrids-off --gcf-cutoffs 0.3,0.7 -m 4.0”.

### Web implementation

The ExMODE platform was developed using a Vue.js (v2.6.14) front-end and a Django (v3.2.15)/Python (v3.10.11) back-end framework. Metadata for publications and datasets are stored in PostgreSQL (v13.5). We used Elasticsearch (v7.16.2) for the resource search engine and Redis (v6.0.16) as an in-memory cache for data management. Task queues are managed using RabbitMQ (v3.8.13), and Nginx (v1.22.1) functions as the reverse proxy. The platform is compatible with major web browsers, including Google Chrome, Opera, Safari, and Firefox.

## Supporting information

Supplemental Table 1, and wil be used for the link to the file on the preprint site.

Supplemental Table 2, and wil be used for the link to the file on the preprint site.

## AUTHOR CONTRIBUTIONS

Denghui Li: Conceptualization; Data curation; writing—original draft; project administration; writing—review and editing. Kailong Ma: Conceptualization; project administration; writing—review and editing. Yulu Zhang: software; Data curation; formal analysis; writing—review and editing. Jieni Wang: Investigation; Methodology; formal analysis; writing—review and editing.

Zhen Cui: Data curation; writing—review and editing. Xiaoqiang Li: formal analysis. Weiwen Wang: Data curation; writing—review and editing. Jiawei Tong: Data curation; visualization. Yang Guo: Conceptualization; writing—original draft. Zongan Wang: Conceptualization; writing—original draft. Peiyun Zeng: software; Data curation; formal analysis. Jieyu Wang: writing—review and editing. Xiaokai Xu: visualization. Nannan Zhang: Data curation. Yingying Zhang: Data curation. Jing Chen: formal analysis. Qingjiang Hu: visualization. Wenzhen Yang: visualization. Zhiyong Li: visualization. Tao Yang: project administration. Wensi Du: Data curation. Zhicheng Xu: Data curation. Zhen Yue: Supervision. Jian Wang: Supervision. Guangyi Fan: Supervision; writing—review and editing. Weijia Zhang: resources; writing—review and editing. Xun Xu: Supervision; writing—review and editing. Liujie Huo: Supervision; writing—review and editing. Xiaofeng Wei: Supervision; writing—review and editing. Liang Meng: Conceptualization; Supervision; resources; writing —review and editing. Shanshan Liu: Conceptualization; supervision; writing—review and editing.

## ACKNOWLEDGMENTS

This work was financially supported by the National Science and Technology Major Project of China (2024ZD1000404). This study was also supported by the High-performance Computing Platform of YZBSTCACC (YaZhou Bay Science and Technology City Advanced Computing Center).

## DATA AVAILABILITY STATEMENT

The datasets and scripts have been deposited on the ExMODE website (https://db.genomics.cn/exmode/) and GitHub (https://github.com/ylzhang1/exmode).

## SUPPORTING INFORMATION

Additional supporting information can be found online in the Supporting Information section at the end of this article.

---

**Figure S1.**
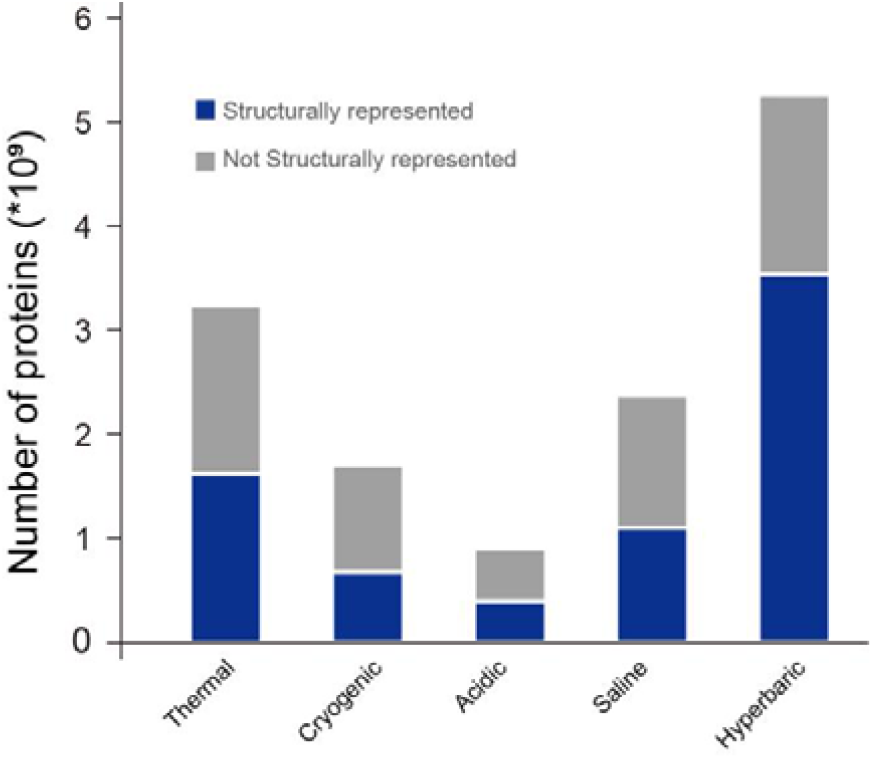
Structural representation of unigenes in different extreme environments

Table S1. The metadata of all samples in ExMODE

Table S2. Genomic information of BGC-rich MAGs

